# Quantitative, image-based phenotyping methods provide insight into spatial and temporal dimensions of plant disease

**DOI:** 10.1101/064980

**Authors:** Andrew M. Mutka, Sarah J. Fentress, Joel W. Sher, Jeffrey C. Berry, Chelsea Pretz, Dmitri A. Nusinow, Rebecca Bart

## Abstract

Plant disease symptoms exhibit complex spatial and temporal patterns that are challenging to quantify. Image-based phenotyping approaches enable multi-dimensional characterization of host-microbe interactions and are well suited to capture spatial and temporal data that are key to understanding disease progression. We applied image-based methods to investigate cassava bacterial blight, which is caused by the pathogen *Xanthomonas axonopodis* pv. *manihotis* (*Xam*). We generated *Xam* strains in which individual predicted type III effector (T3E) genes were mutated and applied multiple imaging approaches to investigate the role of these proteins in bacterial virulence. Specifically, we quantified bacterial populations, water-soaking disease symptoms, and pathogen spread from the site of inoculation over time for strains with mutations in *avrBs2*, *xopX*, and *xopK* as compared to wild-type *Xam*. Δ*avrBs2* and Δ*xopX* both showed reduced growth in planta and delayed spread through the vasculature system of cassava. Δ*avrBs2* exhibited reduced water-soaking symptoms at the site of inoculation. In contrast, Δ*xopK* exhibited enhanced induction of disease symptoms at the site of inoculation but reduced spread through the vasculature. Our results highlight the importance of adopting a multi-pronged approach to plant disease phenotyping to more fully understand the roles of T3Es in virulence. Finally, we demonstrate that the approaches used in this study can be extended to many host-microbe systems and increase the dimensions of phenotype that can be explored.

**Summary:** Novel, image-based phenotyping methods enhance characterization of plant-pathogen interactions.

## Introduction

Plant diseases are responsible for significant reductions in agricultural productivity worldwide, and for many diseases, control strategies are not available (Chakraborty and Newton, 2011). Elucidating the molecular mechanisms that govern host-microbe interactions has a robust track record of leading to the development of new and effective resistance strategies. For example, plant innate immunity employs several tiers of receptors that, at least in some instances, can be transferred among species (Tai et al., 1999; Zhao et al., 2005; Lacombe et al., 2010; Tripathi et al., 2014; Schoonbeek et al., 2015). Similarly, molecular dissection of the mechanisms used by pathogens to induce susceptibility has led to the development of biotechnology methods for blocking pathogenesis (Li et al., 2012; Strauß et al., 2012; Boch et al., 2014). A more complete understanding of the mechanisms used by plant pathogens to cause disease is likely to lead to the development of additional strategies with potential for translation to the field.

Collectively, research conducted over the past few decades has revealed a complicated web of crosstalk that forms our current multi-tiered model of plant-pathogen interactions. Plant pattern recognition receptors (PRRs) initiate immune responses after recognition of conserved microbial features, such as flagellin and EF-Tu for bacteria and chitin for fungi (Macho and Zipfel, 2014). Successful pathogens have evolved effector proteins to suppress defenses and induce susceptibility within their hosts (Win et al., 2012). Resistant hosts may recognize these effectors or their action to trigger robust immune responses (Stam et al., 2014; Khan et al., 2016). Type III effectors (T3Es) secreted into host cells by Gram-negative bacteria are among the most intensely studied pathogen effector proteins, and yet, the function of most T3Es remains unknown.Members of the *Xanthomonas* and *Pseudomonas* genera are among the most common bacterial disease-causing agents and are known to have large and variable effector repertoires (White et al., 2009; Lindeberg et al., 2012; Schornack et al., 2013). Efforts to deduce the role of individual T3Es in bacterial virulence through characterization of effector knockouts have concluded that while collectively important, many individual effectors do not contribute substantially to virulence (Castañeda et al., 2005; Kvitko et al., 2009; Cunnac et al., 2011; Dunger et al., 2012; Xie et al., 2012).

Advances in DNA sequencing technologies have provided a wealth of genomic resources for bacterial species. Using genomics data generated from pathogenic bacteria, we are able to predict T3E repertoires, and the function of individual effectors can then be investigated with genetic knockouts (Baltrus et al., 2011; Bart et al., 2012; Roux et al., 2015; Wei et al., 2015; Teper et al., 2016). Traditional plant disease phenotyping methods have relied on visual assessment of symptoms (Bock et al., 2010) and quantification of pathogen growth in host tissue (Whalen et al., 1991; Tornero and Dangl, 2001; Liu et al., 2015). Visual inspection and scoring of symptoms are likely to be translatable to disease progression within field settings. Inspection and scoring, however, are subject to surveyor bias and may not capture subtle differences in disease severity (Poland and Nelson, 2011). Quantification of pathogen growth is a tractable system for comparison, but fails to provide information regarding the complex spatial patterns of diseases that progress over time. Thus, genetic studies of T3E mutants have likely missed phenotypes that are difficult to measure with traditional methods, and new approaches are needed for exploring additional dimensions of disease phenotypes.

High-throughput, image-based phenotyping methods are revolutionizing many areas of plant biology research (Furbank and Tester, 2011; Fiorani and Schurr, 2013; Araus and Cairns, 2014; Granier and Vile, 2014; Fahlgren et al., 2015; Zaman-Allah et al., 2015). Analysis of plant phenotypes, such as size, shape, color, growth, and leaf area altered by herbivory, can be automatically extracted from image data to observe how such traits change over time (Green et al., 2012; Lamari, 2008). Image-based methods are well suited to characterize the spatial and temporal dimensions of disease symptoms and have been applied to several host-pathogen systems (Mahlein et al., 2012; Rousseau et al., 2013; Bauriegel and Herppich, 2014; Baranowski et al., 2015; Li et al., 2015; Raza et al., 2015). These studies illustrate the range of imaging data that can be generated to automate and quantify detection of disease symptoms. Additionally, these studies emphasize that each imaging assay must be calibrated to detect the critical aspects of the pathosystem being studied.

Cassava (*Manihot esculenta*) is a major staple crop for an estimated 800 million people in Africa, South America, and Asia (FAO, 2013) and is prized as a drought-tolerant plant that is able to thrive on marginal lands. Among the diseases that impact this crop is cassava bacterial blight (CBB), which is caused by the Gram-negative bacterial pathogen *Xanthomonas axonopodis* pv. *manihotis* (*Xam*). CBB disease symptoms are characterized at early stages by water-soaked lesions on leaves and at later stages of infection by wilting and defoliation (Lozano, 1986). Currently, no disease resistance genes have been demonstrated to be effective against CBB, and chemical methods are not an economically feasible form of control for smallholder farmers. Consequently, novel, genetically encoded methods of achieving plant immunity to CBB must be developed.T3Es that are both conserved in *Xam* populations and important for virulence represent 194 possible targets for engineering durable resistance. Previously, the T3E repertoires for 65 *Xam* strains were predicted based on their genomic sequences (Bart et al., 2012). This study predicted 13 to 23 effectors in each strain based on homology to proteins from other systems, excluding the transcriptional activator-like (TAL) effectors that are not resolved by Illumina short read technology. Currently, only a few TAL effectors have been functionally characterized in *Xam* (Castiblanco et al., 2013; Cohn et al., 2014).Notably, several recent studies have explored the use of alternate assays to elucidate the role of T3Es in pathogen virulence (Castiblanco et al., 2013; Cernadas et al., 2014; Cohn et al., 2014).

For this study, we focused on homologs of two previously characterized T3Es, methods such as visual monitoring of symptom development and quantifying bacterial AvrBs2 and XopX, and one predicted T3E, XopK, whose role in virulence is unclear.AvrBs2 contains a glycerol phosphodiesterase domain that is required for its virulence functions in other *Xanthomonas* pathovars (Kearney and Staskawicz, 1990; Tai et al., 1999; Zhao et al., 2011; Li et al., 2015). XopX is involved in suppressing pathogen-triggered immunity (Metz et al., 2005; Sinha et al., 2013; Stork et al., 2015). XopK was first identified in the rice pathogen *Xanthomonas oryzae* pv. *oryzae* (*Xoo*) (Furutani et al., 2006). Previous data indicated that XopK is secreted through the type III secretion system (T3SS) into host cells (Furutani et al., 2009; Schulze et al., 2012). Although the *xopK* gene is conserved in many *Xanthomonas* species, its role in virulence is unknown (Bogdanove et al., 2011; Potnis et al., 2011; Bart et al., 2012; Schulze et al., 2012; Arrieta-Ortiz et al., 2013; Jalan et al., 2013).

Our efforts to phenotypically characterize these *Xam* mutants began with standard methods such as visual monitoring of symptom development and quantifying bacterial growth *in planta* over the course of several days. These methods were limited in resolution, consistency between experiments, and robustness. Consequently, we sought to apply image-based phenotyping methods to study this pathosystem. The first method addresses issues of human bias during scoring by using a low-cost, Raspberry Pi computer and camera to capture and quantify infection over time. The second approach leverages a bioluminescent reporter system within the bacteria to non-invasively monitor pathogen spread throughout the plant (Meyer et al., 2005; Xu et al., 2010). The combination of these methods achieved unprecedented resolution and sensitivity for quantifying the spatial and temporal dynamics of disease progression within the laboratory setting. The results of this study highlight the importance of adopting a holistic approach to phenotyping plant-pathogen interactions to reveal biological functions of virulence proteins and inform development of disease control strategies.

## Results

### *Pathogen growth levels and symptom development for* Xam *type III effector mutants*

To investigate the virulence functions of predicted *Xam* T3Es, we generated *mutations in genes homologous to avrBs2*, *xopX*, and *xopK*. Full gene deletions were created by homologous recombination and initial analysis of pathogen growth levels and symptom development were performed using standard methods (see Materials and Methods) (Whalen et al., 1991; Liu et al., 2015). When inoculated by syringe infiltration in a cassava leaf, wild-type *Xam* populations increased to high levels and caused dark water-soaking symptoms that developed within days of inoculation (Fig. 1). In contrast, mutation of the *hrpF* gene, which encodes a translocon protein required for type III secretion (Buttner et al., 2002), caused a 2-log reduction in pathogen growth levels (Fig. 1, A-B) relative to the wild-type strain and eliminated water-soaking disease symptoms (Fig. 1C). For the Δ*avrBs2* and Δ*xopX* mutants, we observed slightly decreased growth levels compared with wild-type over a total of four independent experiments (Fig. 1A, Supplemental Fig. S1). A generalized linear mixed model (GLMM) adjusted by experiment and technical replicates indicated significant differences in pathogen growth at 5 days post inoculation (dpi) for the Δ*hrpF* (p < 0.001, α = 0.05), Δ*avrBs2* (p < 0.001,α = 0.05), and Δ*xopX* mutants (p = 0.0155, α = 0.05), relative to wild-type. At 6 dpi, the Δ*avrBs2* mutant exhibited a reduction in disease symptoms (Fig. 1C). There were no obvious differences, however, in disease symptom production between the Δ *xopX* mutant and wild-type at 6 dpi (Fig. 1C).

**Figure 1.**
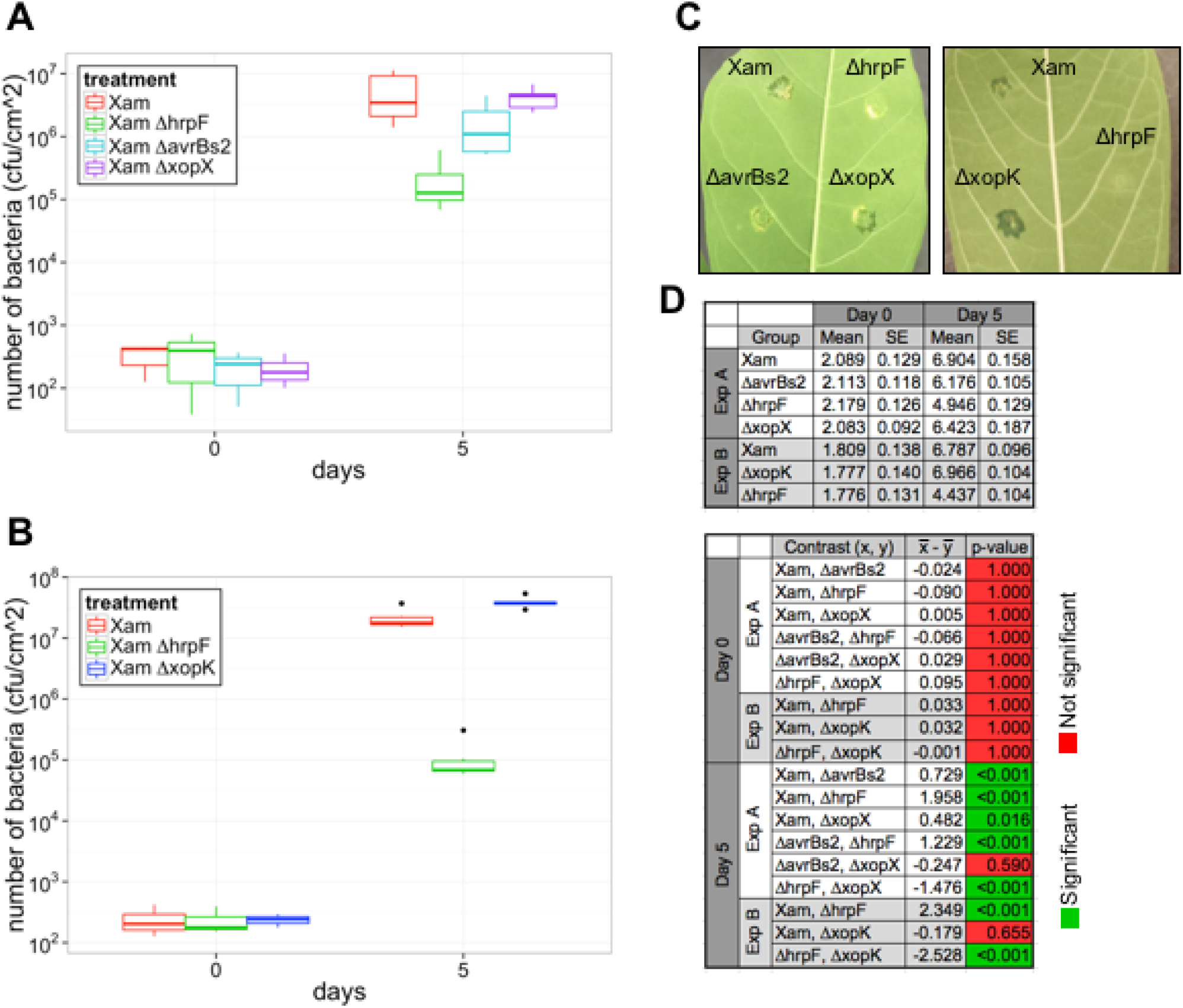
Disease symptom development and growth levels for *Xam* type III effector (T3E) mutants in cassava. A-B, Growth levels of *Xam* wild-type, Δ*hrpF*, Δ*avrBs2*, Δ*xopX*, and Δ*xopK* mutants following syringe infiltration in leaves (OD_600_ = 0.0001). Median, first and third quartiles are shown. The whiskers extend to the highest and lowest data point falling within 1.5∗IQR (inter-quartile range). Dots represent outliers that fall outside 1.5∗IQR. Each experiment was repeated three additional times (see Supplemental Fig. S1). C, Comparison of disease symptoms caused by T3E mutants on leaves at 6 days after syringe infiltration (OD_600_ = 0.0001). D, Results of generalized linear mixed model analysis, combining bacterial growth data from all replicate experiments. Combined estimated means and standard error (SE) are presented, as well as the difference between the means and the p-values for each pairwise statistical contrast.

In contrast to the Δ*avrBs2* and Δ*xopX* mutants, we observed that the Δ*xopK* mutant exhibited pathogen growth levels that were either similar to or elevated compared to wild-type across four experiments (Fig. 1B, Supplemental Fig. S1). A GLMM indicated the growth differences observed for the Δ*xopK* mutant at 5 dpi were not significantly different from wild-type (p = 0.655, α = 0.05). However, water-soaking disease symptoms caused by the Δ*xopK* mutant appeared enhanced at 6 dpi (Fig. 1C). To address the limitations of visual assessment of disease and to better understand the temporal dimension of this phenotype, we sought to develop a quantitative method of assessing symptom development with increased time resolution.

### Image-based quantification of disease symptom development

To develop a low-cost imaging system for semi-automated quantification of disease symptom development, we used Raspberry Pi microcomputers and camera boards to image the abaxial side of the leaf during symptom development (Fig. 2A). Following 265 data collection, regions of interest (ROI) were selected manually from the image stacks that contained each inoculated spot, and image analysis was performed using an ImageJ macro script (Fig. 2A, Supplemental Fig. S2).

**Figure 2.**
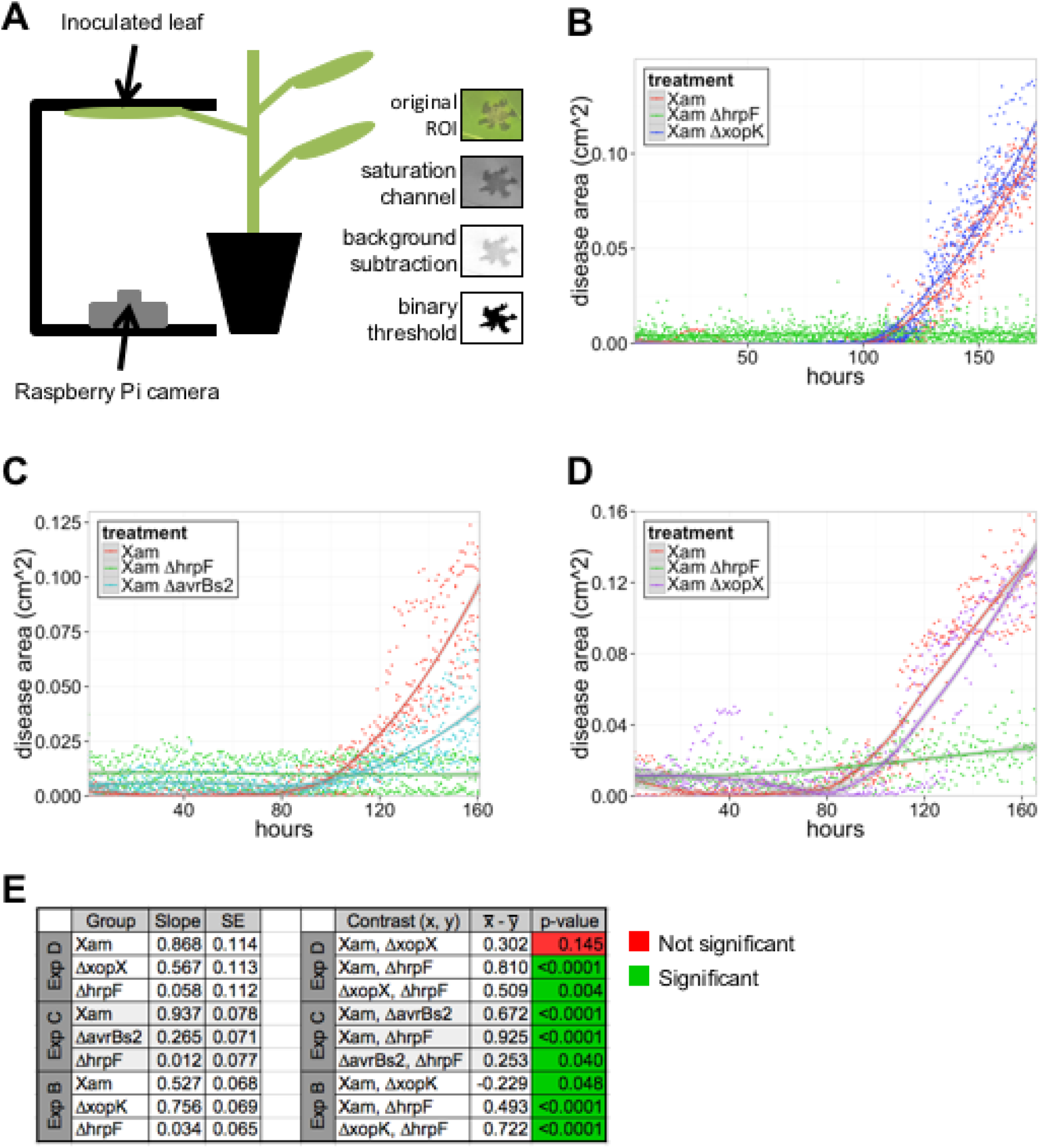
Quantification of disease symptoms caused by *Xam* on cassava leaves using imaging. A, Illustration of the imaging set-up. Leaves were syringe infiltrated with bacterial solutions (OD_600_ = 0.001) and taped to a black surface for imaging of the abaxial side of the leaf. Raspberry Pi microcomputer with attached camera collected images once per hour. Image analysis steps in ImageJ are shown and described in Materials and Methods. The number of black pixels was quantified to determine the area of disease. B D, Quantification of water-soaking symptoms caused by the *Xam* wild-type strain and three mutants over time. Dots represent individual measurements determined from image analysis, and local regression fitted curves are plotted for each bacterial strain. Shaded areas represent the 95% confidence interval for each curve. The experiment was repeated three additional times with similar results (see Supplemental Fig. S3). E, Results of generalized linear mixed model analysis, combining data from all replicate experiments.Combined estimated slopes and standard error (SE) are presented, as well as p values for each pairwise statistical contrast.

We quantified water-soaking disease symptoms caused by wild-type *Xam*, along with the Δ *xopK* and the Δ*hrpF* mutants (Fig. 2B). For the wild-type strain, disease symptoms began appearing at approximately hours post inoculation. As expected, the Δ*hrpF* mutant did not produce disease symptoms throughout the course of the experiment. For the Δ*xopK* mutant, an increased rate of symptom accumulation was observed relative to the wild-type strain in the initial phase of disease appearance over four separate experiments (Fig. 2B, Supplemental Fig. S3). To compare the rates of symptom accumulation for these strains, we performed a GLMM analysis, adjusted by experiment and technical replicates. Additionally, we adjusted for heteroskedasticity by allowing the variance to be exponentially related to time. This analysis indicated that slopes for both the Δ*hrpF* (p < 0.0001, α = 0.05) and Δ*xopK* (p = 0.0476, α = 0.05) mutants were significantly different from wild-type. Thus, despite being a predicted T3E, mutation of the *xopK* gene induces disease symptoms more rapidly during infection than wild-type *Xam*.

To further investigate the impacts of other T3E mutations on disease symptom progression, we performed imaging of cassava leaves inoculated with the *Xam*Δ*avrBs2* and Δ*xopX* mutants (Fig. 2C, D). As with the experiments involving the Δ*xopK* mutant, we used GLMM, adjusted by experiment, technical replicates, and heteroskedasticity. The Δ*avrBs2* mutant exhibited significantly decreased disease symptom progression relative to wild-type (p < 0.0001, α = 0.05), consistent with visual observations of disease (Fig 1). For the Δ*xopX* mutant, while symptom progression appeared slightly delayed compared to wild-type, this effect was not significantly different from wild-type across all experiments (p = 0.1447, α = 0.05). Thus, in the T3E repertoire of *Xam*, AvrBs2 has a greater contribution to early symptom progression than XopX.

### Characterizing bacterial spread in host tissue

To observe the ability of T3E mutants to spread systemically during infection, we developed a method to track bacterial spread *in planta.* This is particularly important for *Xam* because it spreads through the host vascular system to cause disease (Lozano, 1986). Therefore, bacterial growth and symptom development at the site of inoculation are descriptive of only a small part of pathogenesis. It has been shown that bioluminescent bacterial strains can be used to detect pathogen presence in host tissue (Bogs et al., 1998; Meyer et al., 2005; Xu et al., 2010).

To determine if we could detect *Xam* and visualize spread from an initial infection site *in planta*, we generated bioluminescent strains by introducing a plasmid driving constitutive expression of the LUX operon into *Xam* and the T3E mutants by conjugation. The inoculated areas were imaged in a dark chamber to detect the bioluminescence signal after syringe inoculation of LUX strains into cassava leaves. Local areas infiltrated with wild-type *Xam* exhibited a circular region of bioluminescence that appeared by 4 dpi, followed by spread into surrounding tissues (Fig. 3). This spread extended beyond the 308 visible area of water-soaking symptoms observed at 9 dpi. These results indicate *Xam* proliferates locally at the site of inoculation before invading into the nearby vasculature, and that bacterial spread can be observed in regions of the plant that do not yet exhibit visible symptoms.

**Figure 3.**
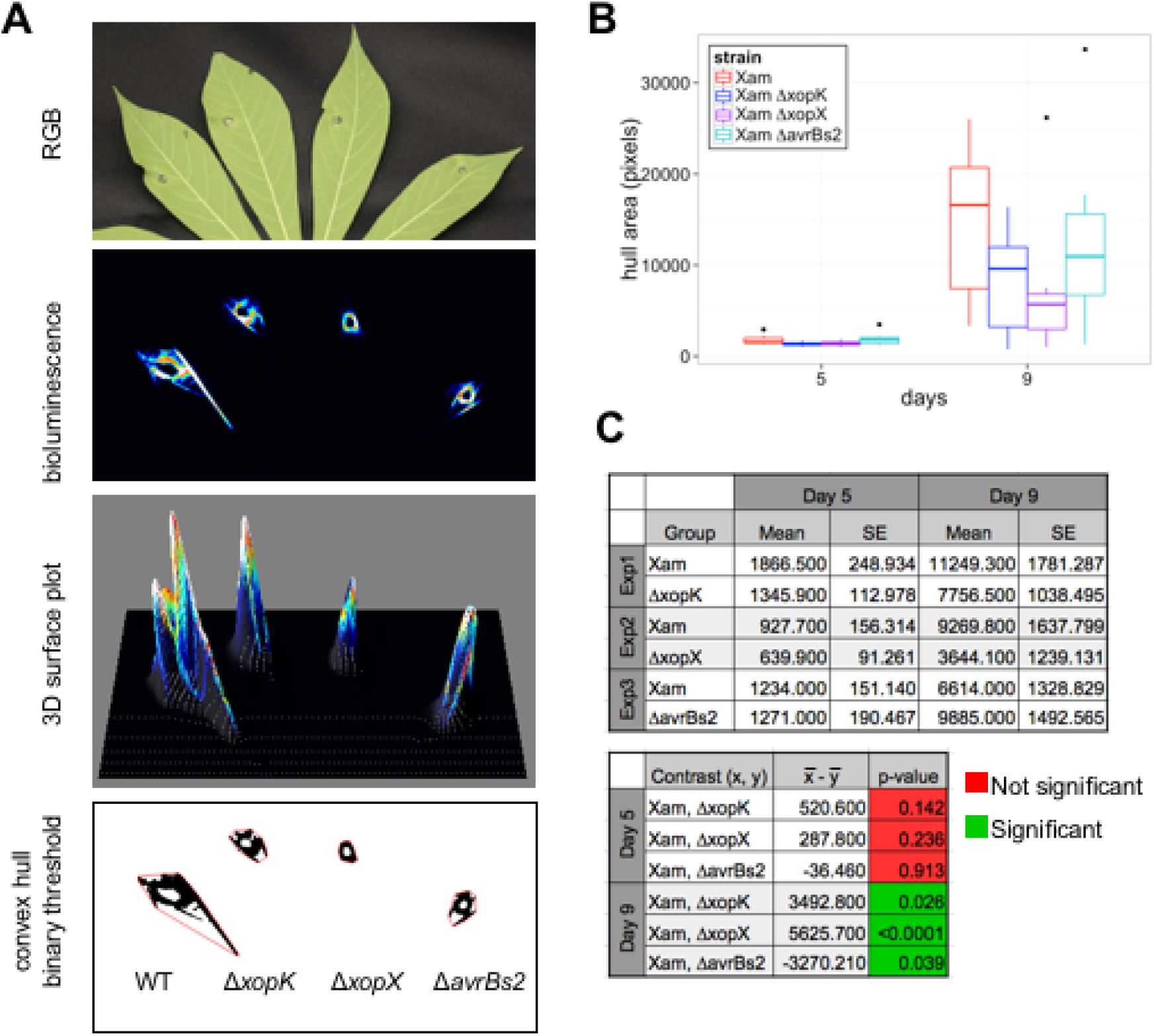
Bioluminescence imaging of *Xam* spread in cassava leaves. A, The bioluminescence reporter pLUX plasmid was introduced into *Xam* strains. Leaves were inoculated with bacterial solutions (OD_600_ = 0.01) using syringe infiltration. RGB images of inoculated leaves reveal symptom development. Bioluminescence was visualized in a dark chamber with a 5-10 min exposure. Image processing was performed with ImageJ to select the area of bioluminescence and the convex hull of the resulting shapes were analyzed. B, Representative quantification of convex hull for wildtype Xam and three mutants. Additional replicate experiments are shown in Supplemental Fig. S4. C, Results of generalized linear mixed model analysis of convex hull area, combining data from all replicate experiments. Combined estimated means and standard error (SE) are presented, as well as the difference between the means and the p-values for each pairwise statistical contrast.

To quantify the changes in bacterial luminescence observed from these images, we applied image analysis methods for measuring both the convex hull area and maximum span across the convex hull, which estimate the total area invaded and the maximum linear distance of spread by the pathogen, respectively. Compared to wild-type *Xam*, the *ΔxopX* mutant exhibited reduced spread, observed in five independent experiments (Fig. 3, Supplemental Fig. S4). A GLMM, adjusted by experiment and technical replicates, indicated at 9 dpi this reduction in spread quantified by convex hull area was statistically significant (p < 0.0001, α = 0.05). We observed significant reduction in pathogen spread for the *ΔavrBs2* (p = 0.0388, α = 0.05) and Δ*xopK* mutants (p = 0.0257, α = 0.05). Further consideration of these data revealed that experimental noise reflected the timing of pathogen entry into the plant vasculature (Fig. 3, Supplemental Fig. S4). These results highlight the importance of invasion into host vasculature by *Xam* for virulence and the power of image-based phenotyping assays for visualizing this dynamic process.

### *Characterizing bacterial spread* in vitro

Given the phenotype observed for reduced spread in host tissue, next we examined if any of the T3E mutants affected motility *in vitro* (Supplemental Fig. S5).Bacteria were plated at the center of soft agar plates, which allowed them to spread across the surface of the media (Lee et al., 2003; Tian et al., 2015). Using a Raspberry Pi-controlled camera, images of the plates were taken for several days, and the area of bacterial spread was quantified (Supplemental Fig. S5). We observed that the *ΔhrpF, ΔavrBs2*, *ΔxopX*, and *ΔxopK* mutants do not have any apparent motility defects on soft agar. These results suggest that observed defects in mutant movement through host vascular tissue are due to factors other than intrinsic bacterial motility.

### Imaging bacterial colonization in other pathosystems

Since tracking bacterial colonization with a bioluminescent reporter was successful in cassava, we wanted to expand the use of this method to other pathosystems. To explore this, we generated bioluminescent *Xanthomonas campestris* pv. *campestris* (*Xcc*) and *Xanthomonas euvesicatoria* (*Xe*) by conjugation. Following syringe inoculation of bioluminescent *Xam* and *Xcc*, bacteria can be visualized near the site of infection and, in the case of *Xam,* at distal sites following spread into the vasculature (Fig. 4). Since *Xe* infection in pepper and tomato results in leaf spot disease, dip inoculations were used for this strain to mimic field infections. Bioluminescent *Xe* could be readily detected in distinct puncta across the leaf following dip inoculation (Fig. 4). These results demonstrate the utility of bioluminescent technologies to track pathogens in diverse crop plants.

**Figure 4.**
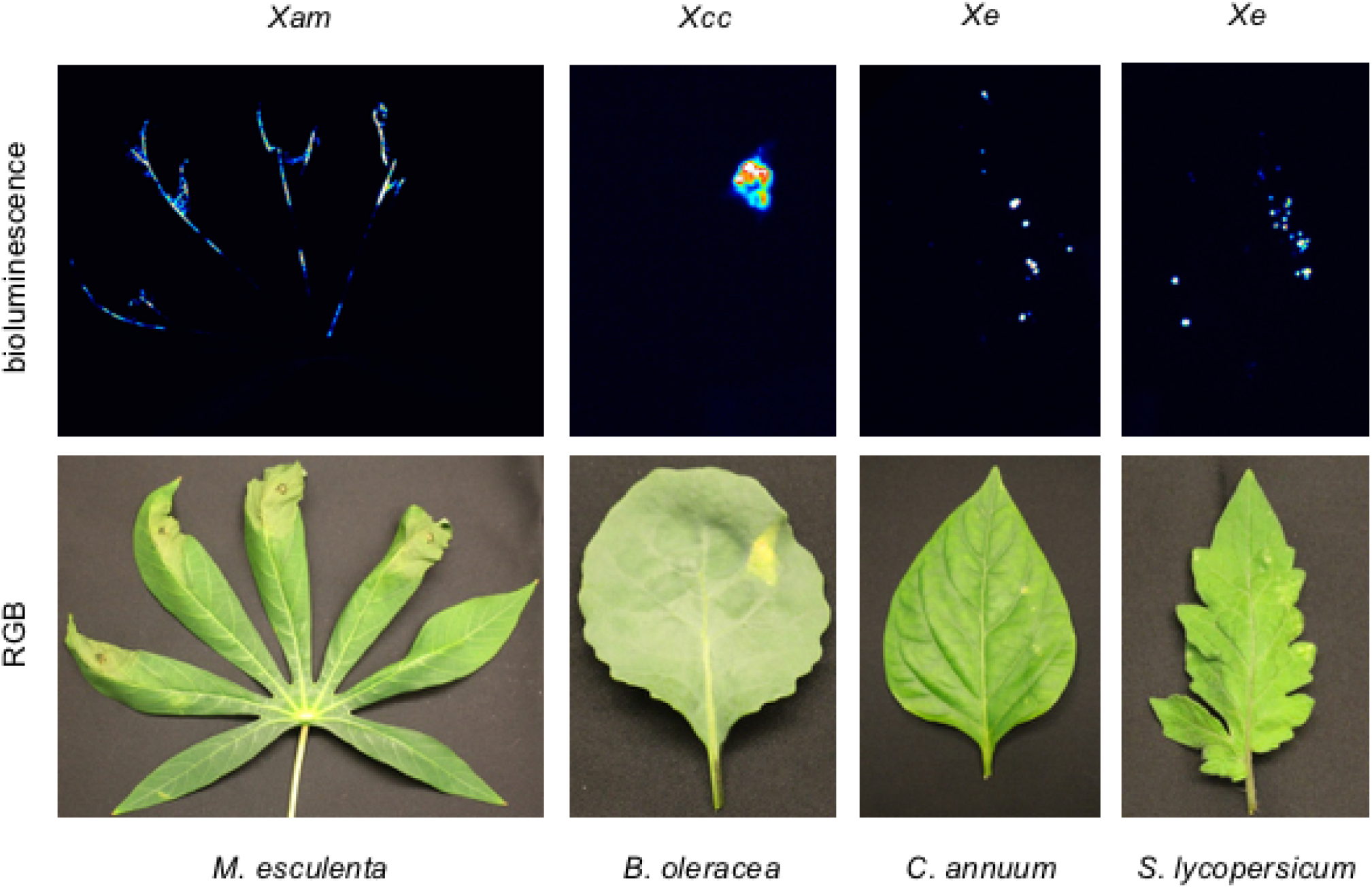
Comparison of vascular and leaf spot pathogens. *Xam* and *Xcc* were visualized 615 following syringe inoculation (OD_600_ = 0.01) at 18 days post inoculation (dpi) and 12 dpi, respectively. Bacterial spot on pepper and tomato were visualized following dip inoculation (OD_600_ = 0.5) of *Xe* at 9 dpi.

## Discussion

In complex interactions between hosts and pathogens, T3Es are key targets for resistance strategies. The specific virulence functions of T3Es, however, can be difficult to determine, and their characterization remains a primary goal of researchers within the plant-microbe interaction field. Traditionally, relative virulence of pathogen strains is quantified through visual assessment of plant symptoms and destructive harvesting to measure pathogen populations over time. However, these methods alone often do not provide enough information to make conclusions about T3E functions during infection.Furthermore, CBB has a number of challenges for experimental study. For example, *Xam* is able to spread *in planta* through both mesophyll and vascular tissue (Boher and Verdier, 1994). The pattern of colonization for these distinct tissue types likely contributes to observed experimental noise. Furthermore, cassava plants are propagated from vegetative cuttings, so it is difficult to obtain plants that are developmentally synchronized and physiological aspects of cassava plants such as latex content of leaves may vary from one experiment to another. Finally, it is notable that cassava is a field grown crop with a long generation time (12 months) and development of phenotyping methods from which observations will translate to field setting is desirable.

To address these limitations, we applied image-based phenotyping methods that enable quantification of spatial and temporal dynamics for plant-pathogen interactions (Figure 5). Measuring symptom area with increased time resolution, enabled by automated imaging, allowed us to observe an enhanced rate of early disease accumulation for the Δ *xopK* mutant. This phenotype demonstrates the importance of a pathogen’s ability to establish infection for overall virulence. While XopK is a predicted T3E effector that carries the hallmarks of a secreted virulence factor (Furutani et al., 2006; Furutani et al., 2009; Schulze et al., 2012), its roles during infection appear to be complex. The XopK protein sequence contains 54% hydrophobic residues and several predicted transmembrane domains. Thus, it is possible this protein is associated with host cell membranes following secretion. Our observations for XopK are contrasted with the roles of AvrBs2 and XopX, which exhibit reduced virulence phenotypes, consistent with 380 previous studies (Kearney and Staskawicz, 1990; Metz et al., 2005; Zhao et al., 2011; Sinha et al., 2013; Li et al., 2015; Stork et al., 2015). Our results further illustrate that T3E mutants may impact certain aspects of host-pathogen interactions more than others.For example, XopX contributes more to *Xam* proliferation in the host than to disease symptom progression, while AvrBs2 contributes significantly to both aspects of infection.Thus, combining data from several phenotyping approaches is necessary for determining 386 the roles of T3Es in virulence (Table 1).

**Figure 5.**
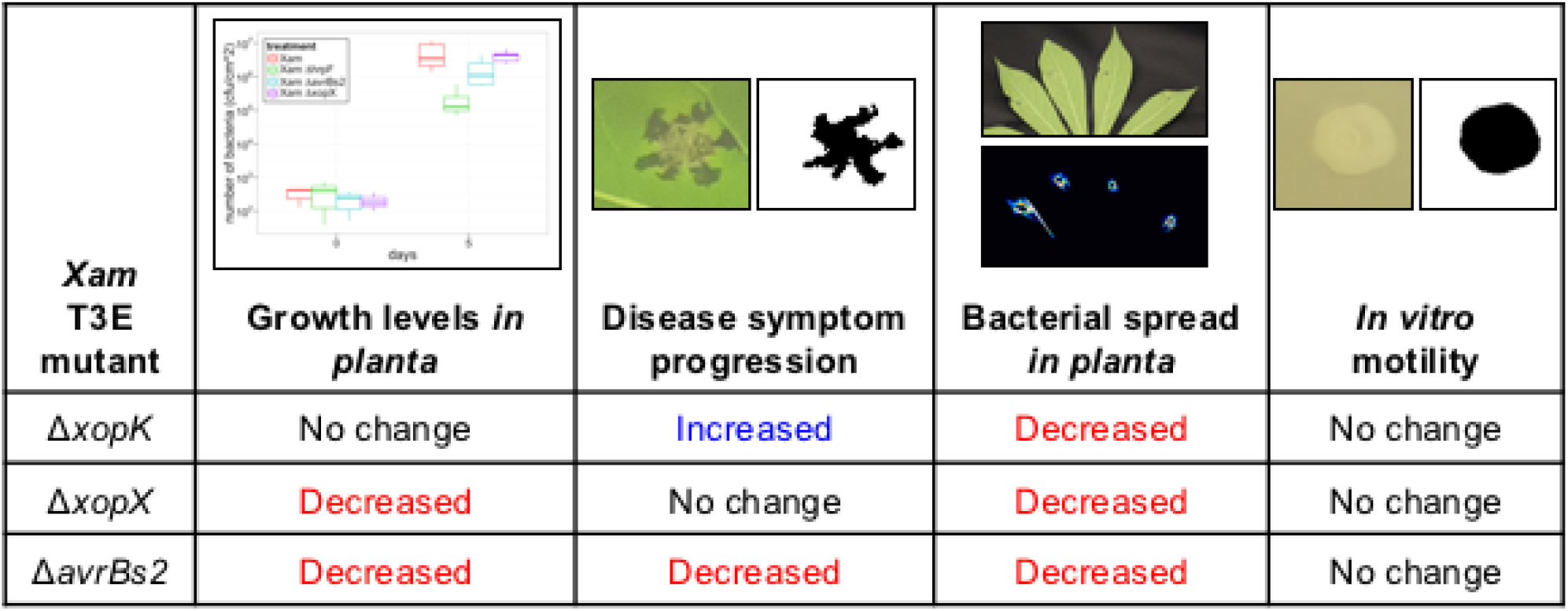
Summary for type III effector mutant phenotypes as revealed by different 620 phenotyping methods.

Another challenge of studying plant-pathogen interactions is visualizing and quantifying a pathogen’s location and movement in host tissue. As we observed imaging *Xam* strains in cassava, bioluminescence is a powerful tool for studying the spatial and temporal dimensions of pathogen spread in host tissue. Previously, bioluminescent bacterial strains were used to visualize *Xcc* infection in *Arabidopsis thaliana* (Meyer et al., 2005). Also bioluminescent strains of the tomato pathogen *Clavibacter michiganensis* were used to track bacterial movement following grafting and during germination (Xu et al., 2010). Imaging of bioluminescence with increased spatial resolution allows for quantification of patterns of pathogen spread, as we observed for *Xam* infection in cassava leaves. Our studies visualized the process by which *Xam* invades host vasculature. The experimental variation we observed suggests this is a dynamic process impacted by a range of factors, such as proximity to primary and secondary leaf veins, leaf developmental status, and environmental conditions. These variables can be investigated using our imaging approaches. In particular, environmental factors such as humidity and heat are known to be important for CBB severity in the field (Boher and Verdier, 1994). How these conditions promote disease severity is currently unknown but may be related to mechanisms of pathogen spread in host tissue. These factors can be readily explored by direct imaging using an environmentally controlled chamber.

While image-based phenotyping approaches offer great promise, characterizing plant-pathogen interactions with such methods also has many challenges, in part due to the diverse range of symptoms and multiple scales at which disease occurs. For each pathosystem, one must first identify the relevant aspects of the infection that can be imaged and the relevant metrics that are needed to characterize the system. This initial study relied on visible light imaging. In future studies, other wavelengths of the electromagnetic spectrum that provide additional information for characterizing disease will be considered. Hyperspectral imaging, which collects spectral data for every pixel, has been used to classify and quantify several diseases that infect sugar beet (Mahlein et al., 2012; Leucker et al., 2016). Thermal imaging offers another approach for detecting disease in plant canopies, since many diseases impact transpiration rates and, therefore, plant surface temperature. While environmental variation can introduce challenges into thermal imaging, Raza et al. addressed this issue by using a combination of thermal and visible light imaging, along with depth estimation, to detect diseased tomato plants using a machine learning approach (Raza et al., 2015). In a laboratory context, fine-scale imaging of disease symptoms on a single leaf may be the optimal approach for investigating certain experimental questions, while whole-plant or field-scale imaging would be necessary for detecting disease in an agricultural context. Unmanned aerial vehicles performing multispectral imaging were used at the field scale to examine abiotic stress in maize (Zaman-Allah et al., 2015). Biotic stresses likely could be detected using similar signatures from aerial imaging.

This study presents a new approach to investigation of plant-pathogen interactions. Inspired by the advances of image-based phenotyping methods in other areas of plant biology, we have applied similar methods to quantifying spatial and temporal dimensions of disease development. This approach expands the potential range of phenotypes that can be explored, enabling insights that are difficult to obtain by traditional methods. Since many smallholder farmers in the developing world rely on cassava for food security, low-cost monitoring devices that enable rapid detection of disease outbreaks in the field would be beneficial. In an ideal scenario, monitoring devices would be deployed to cassava fields in disease-prone regions and transmit data over wireless networks to give farmers early warning of disease outbreaks. Many technical and logistical challenges would need to be overcome to achieve this goal.However, our image-based methods for detecting disease represent a first step in developing capabilities for such a device. Remote detection of disease through imaging or other means is a promising approach for plant pathology research that can be translated from the laboratory to the field. While characterizing each plant-pathogen system has its own unique challenges, image-based phenotyping methods can be adapted for many systems and offer the potential to revolutionize plant disease identification and quantification.

## Materials and methods

### Bacterial strains and plant varieties

The following strains of *Xanthomonas axonopodis* pv. *manihotis* (*Xam*) were used:*Xam668*, *Xam668 ΔavrBs2*, *Xam668 ΔhrpF*, *Xam668 ΔxopK*, and *Xam668 ΔxopX*. For experiments performed in cassava (*Manihot esculenta*), variety 60444 was used, except in experiments noted were variety TME7 was used. Strains used for other pathosystems include: *Xanthomonas euvesicatoria* (*Xe* 85-10) and *Xanthomonas campestris* pv. *campestris* (*Xcc* 8004). For experiments in pepper, *Capsicum annuum* variety ECW was used. For experiments in tomato, *Solanum lycopersicum* variety M82 was used. For experiments in broccoli, *Brassica oleracea*, was used.

### Generation of bacterial mutants

To generate gene deletion mutants in bacteria, we used a homologous recombination strategy, in which genomic regions flanking each gene of interest were first cloned into the pENTR-d-TOPO vector (Invitrogen). Blunt-end cloning of the PCR products was performed to introduce the PCR fragments into pENTR-d-TOPO.Restriction sites added with the primers were used to create a version of the vector with 5' and 3' flanking regions adjacent to each other. Gateway cloning was used to transfer this cassette into the pLVC18-sacBR vector, which enabled a sucrose counter-selection approach to be used in creating unmarked gene deletions (Logue et al., 2009). This vector was conjugated into *Xam* using a triparental mating system, with an *E. coli* helper strain containing the pRK600 plasmid. Mutants with the unmarked gene knockout were confirmed with PCR.

### *Bacterial inoculations and growth monitoring* in planta

To quantify bacterial growth in plant tissue, leaves were inoculated with solutions of bacterial strains (OD_600_ = 0.0001) re-suspended in 10 mM MgCl_2_. Prior to inoculation, the leaf was wounded with a razor blade, and the bacterial solutions were injected into the leaf with a needleless 1-mL syringe. Approximately 0.1 mL of bacterial solution was injected at each inoculation site.

For each time point that bacterial growth was monitored, leaf punches were taken from inoculated regions. A 1-cm diameter cork borer was used to excise a leaf disk, which was then ground with a tungsten carbide bead in 200 μL of 10 mM MgCl_2_ using a TissueLyser (Qiagen). Dilutions of the leaf lysate were then plated on NYG agar media with 100 μg μL^-1^rifampicin, and *Xam* colonies were counted to estimate the number of bacteria in the original leaf sample.

### Image acquisition and analysis for quantification of disease symptoms

Leaves were inoculated with solutions of bacterial strains (OD_600_ = 0.001). The leaves were then taped to a black surface, such that the abaxial side of the leaf faced downward toward a camera. Images of disease symptoms were taken with Raspberry Pi 5MP camera boards controlled by Raspberry Pi Model B microcomputers. Hourly automated image collection was done with Cron. For all experiments shown, images were taken hourly. Image stacks were analyzed with FIJI, running ImageJ version 2.0.0-rc-30/1.49v (Schindelin et al., 2012; Schindelin et al., 2015). The scale of the image stacks was determined with a ruler included in the image, enabling unit conversion from pixels to cm.

A region of interest (ROI) was selected that contained each inoculated area. An ImageJ macro script was used to select and quantify the area of water-soaking symptoms for each ROI (Supplemental File S1). The ROI stacks were converted to HSB (hue-saturation-brightness) color space. After selecting the saturation channel, a background subtraction was performed. Then, the area of disease was selected by creating a binary image with black pixels representing symptomatic regions and white pixels representing non-symptomatic tissue. The binary images were created by applying a threshold calculated by the following,

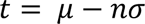

where *t* = threshold, μ = mean saturation value of the ROI stack, and σ = standard 501 deviation of saturation values for the ROI stack. For the variable *n*, a range of values were tested to determine the appropriate threshold for selecting disease symptoms. The area of black pixels in the binary images was then quantified to determine the area of disease symptoms for each ROI.

### Bioluminescence imaging and analysis

Bioluminescent bacteria were generated by conjugating the pLUX plasmid into *Xanthomonas* strains of interest. Leaves were inoculated with solutions of these strains (OD_600_ = 0.01), with one strain inoculated per leaf lobe. Images were taken using a Star I CCD digital camera system (Photometrics Ltd.) or a PIXIS 1024B (Princeton Instruments) that was contained in a blackout chamber. The cameras were operated with WinView/32 software version 2.5.22.0 (Princeton Instruments) or µManager v1.4.22 (Open Imaging), respectively.

Using ImageJ, each image was converted to an 8-bit TIFF file. Then, the Auto Enhance Contrast function was applied, and a binary image was created with the Auto Threshold function using the MaxEntropy algorithm (Kapur et al., 1985). Next, the Despeckle and Invert functions were applied to create an image in which the bioluminescent signal was represented by black pixels. Finally, the Hull and Circle plugin v2.1b (http://rsb.info.nih.gov/ij/plugins/hull-circle.html) was used to select and quantify the convex hull for each shape representing the bioluminescence signal.

### In vitro *bacterial motility assays*

The ability of *Xam* strains to spread *in vitro* was observed on NYG media containing 0.25% agar and 100 μg μL^-1^ rifampicin. 10 μL of a bacterial solution (OD_600_=0.1) was spotted in the center of each plate. A Raspberry Pi camera was set to image the plates from a top-view. Images of the plates were taken over several days, as the bacteria spread from the center of the plates. The ROI containing the bacterial colony on each plate was selected, and an ImageJ macro script was used to create a binary image where black pixels represented the area of the bacterial colony (Supplemental File S2). The number of black pixels was then quantified to determine the bacterial area on the plate.

### Data and statistical analysis

All graphs and data analyses were generated with R version 3.1.3 (R Core Team, 2015), using the following packages: ggplot2 version 1.0.1 (Wickham, 2009), knitr version 1.11 (Xie, 2015), lme4 version 1.1-11 (Bates et al., 2015), lsmeans version 2.20-23 (Lenth, 2015), multcomp version 1.4-4 (Hothorn et al., 2008), nlme version 3.1-128 (Pinheiro et al. 2016), plyr version 1.8.3 (Wickham, 2011), reshape2 version 1.4.1 (Wickham, 2007), and scales version 0.3.0 (Wickham, 2015). For each data set, a generalized linear mixed model was used to assess the responses as a function of different bacterial strains and over time while adjusting for correlation structures due to repeated measures and cross-experimental variation. After the models were fit and checked for underlying assumptions, statistical contrasts were implemented using a post hoc Tukey’s t test to compare the effect of interest across the different bacterial strains using an α level of 0.05 as the cutoff for statistical significance.

### Supplemental material

The following supplemental materials are available.

Supplemental Figure S1. Replicate experiments for analysis of *Xam* T3E mutant growth *in planta* shown in Fig. 1.

Supplemental Figure S2. Image analysis optimization for water-soaking disease symptoms caused by *Xam*.

Supplemental Figure S3. Replicate experiments for analysis of water-soaking symptoms shown in Fig. 2.

Supplemental Figure S4. Replicate experiments for analysis of *Xam* spread in leaves using bioluminescence imaging shown in Fig. 4.

Supplemental Figure S5. *In vitro* motility of *Xam* T3E mutants.

Supplemental File S1. ImageJ macro script used to quantify water-soaking disease symptoms.

Supplemental File S2. ImageJ macro script used to quantify bacterial spread on soft agar plates.

## Author contributions

A.M.M. designed and performed experiments, analyzed data, and co-wrote the paper. S.J.F. created bioluminescent bacterial strains, performed experiments, and edited the paper. J.W.S. and C.P. performed experiments and provided technical assistance. J.C.B.performed statistical analyses. D.A.N. co-supervised the development of the bioluminescent imaging techniques and edited the paper. R.B. designed experiments, supervised the study, and co-wrote the paper.

## Financial sources

This work was supported by the Donald Danforth Plant Science Center

## Acknowledgments

We would like to thank Malia Gehan and Noah Fahlgren for their help with the image acquisition and analysis.

---

**Supplemental Figure S1.**
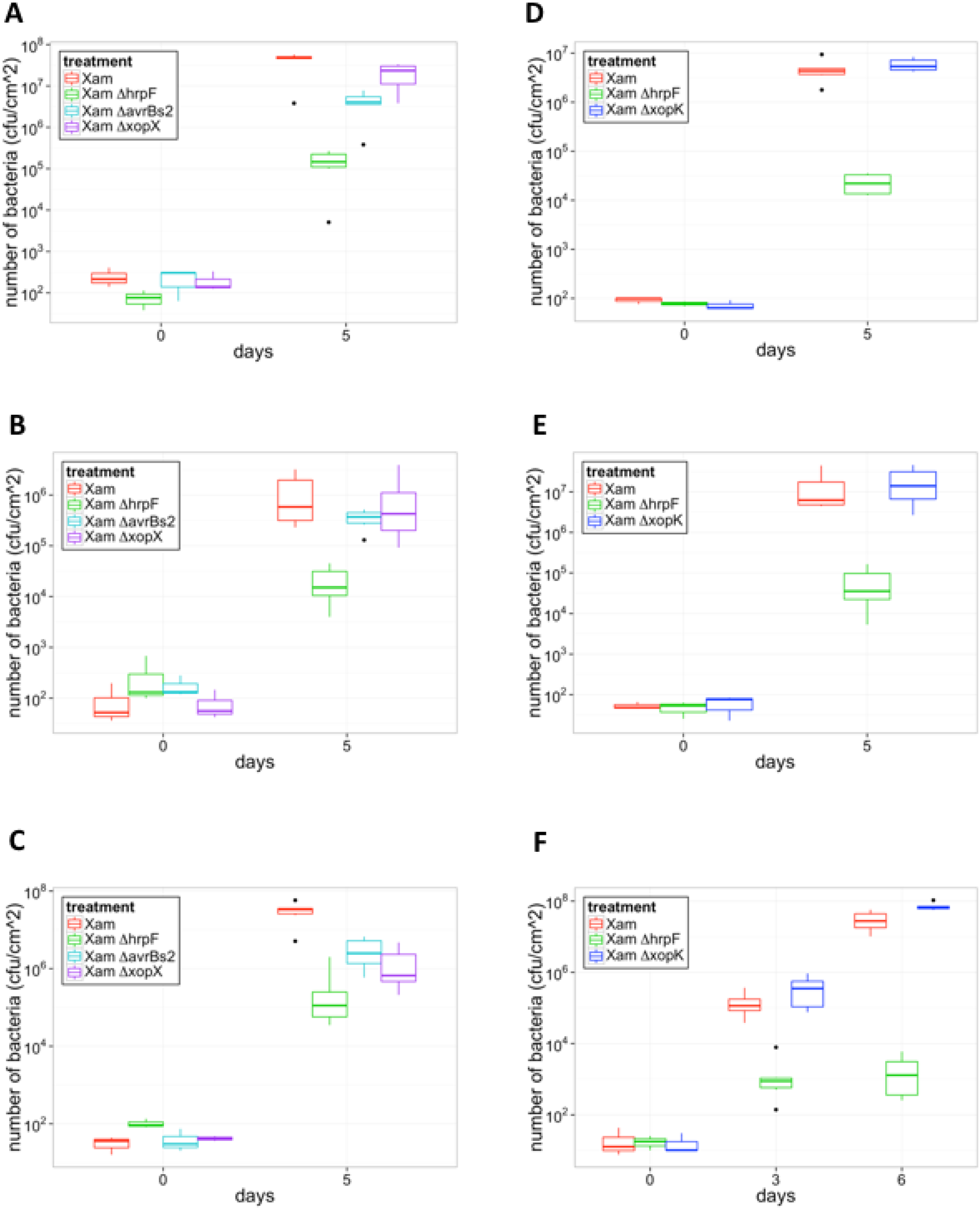
Replicate experiments for analysis of *Xam* T3E mutant growth *in planta* shown in Figure 1. A-C, Three additional experiments to quantify growth levels of the *Xam* wild type strain, and the *Xam*Δ*hrpF*, Δ*avrBs2*, and Δ*xopX* mutants following syringe infiltration in leaves (OD_600_ = 0.0001). D-F, Three additional experiments to quantify growth levels of wild-type *Xam*, and the Δ*hrpF* and Δ*xopK* mutants following syringe infiltration in leaves (OD_600_ = 0.0001).

**Supplemental Figure S2.**
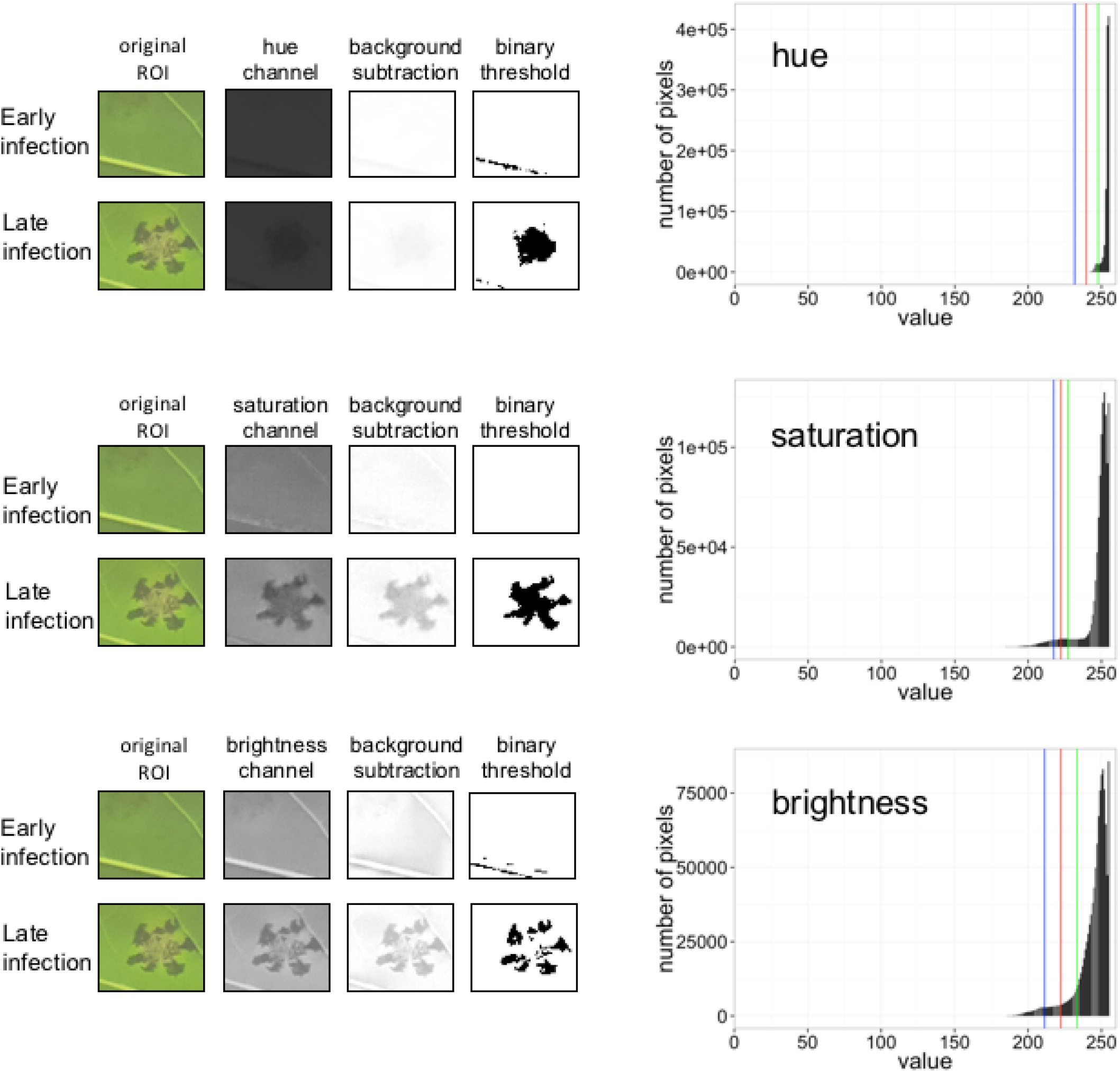
Image analysis optimization for water-soaking disease symptoms caused by *Xam*. Following conversion of regions of interest (ROI) to hue-saturation-brightness (HSB) color space and background subtraction, the distributions of pixel values for each channel were examined. Disease symptoms are best represented by pixel values in the lower tail of the distribution for the saturation channel. A range of threshold values were applied to the images to create binary images, according to the formula t = μ – nσ, where t = threshold, μ = mean saturation value for the image stack, and σ = standard deviation of saturation values for the image stack. Graphs show example thresholds where n = 3 (blue line), n = 2.5 (red line), and n = 2 (green line). The binary images on the right show the results of applying the n = 3 threshold.

**Supplemental Figure S3.**
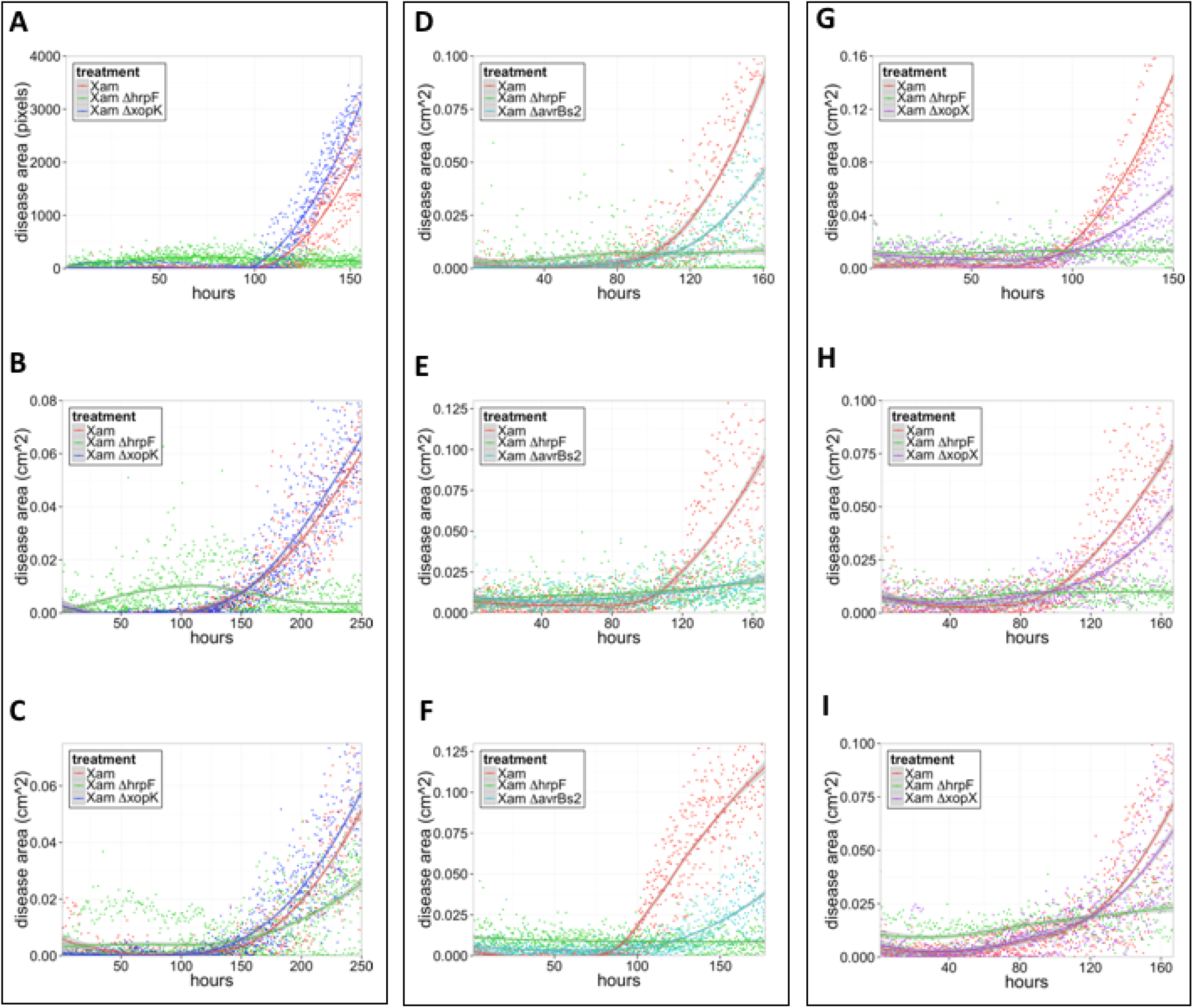
Three additional replicate experiments for analysis of water-soaking symptoms shown in Fig. 2. Dots represent individual measurements determined from image analysis, and local regression fitted curves are plotted for each bacterial strain. Shaded areas represent the 95% confidence interval for each curve. For the experiment shown in A, the image distances were not scaled to cm, so values are reported in pixels. Experiments shown in panels D-F and H-I were performed with cassava variety TME7.

**Supplemental Figure S4.**
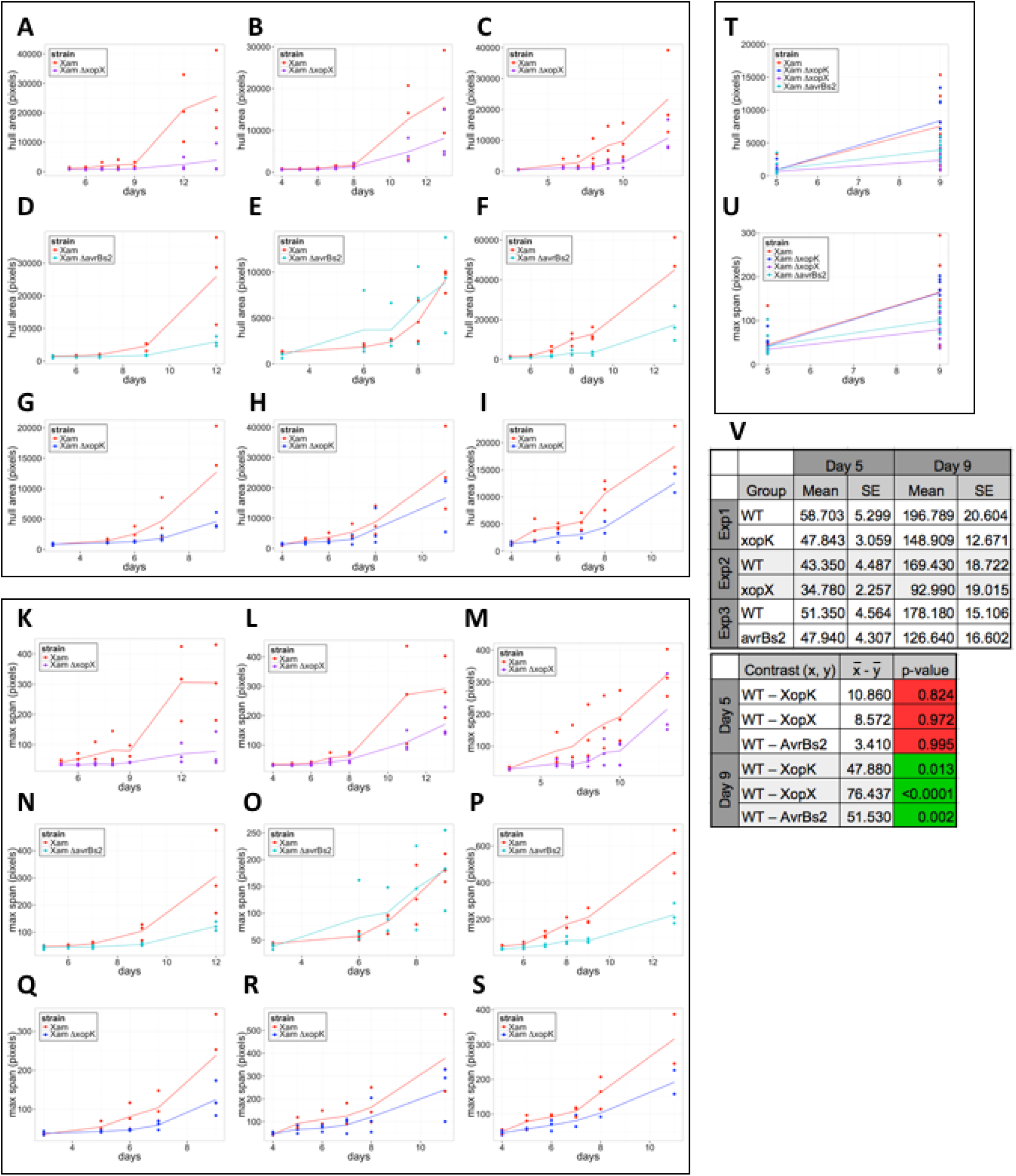
Replicate experiments for analysis of *Xam* spread in leaves 649 using bioluminescence imaging shown in Fig. 3. Three additional experiments quantifying the convex hull area for the *Xam*Δ*xopX* (A-C), Δ *avrBs2* (D-F), and Δ *xopK* (G-I) mutants. Quantification of the maximum span across the convex hull for the same experiments (K-S). Quantification of the convex hull area (T) and maximum span (U) for a fourth replicate experiment including the three mutants. Dots represent individual measurements; lines connect the mean at each time point. V, Results of generalized linear mixed model analysis for maximum span results, combining data from all replicate experiments. Combined estimated means and standard error (SE) are presented, as well as p values for each pairwise statistical contrast.

**Supplemental Figure S5.**
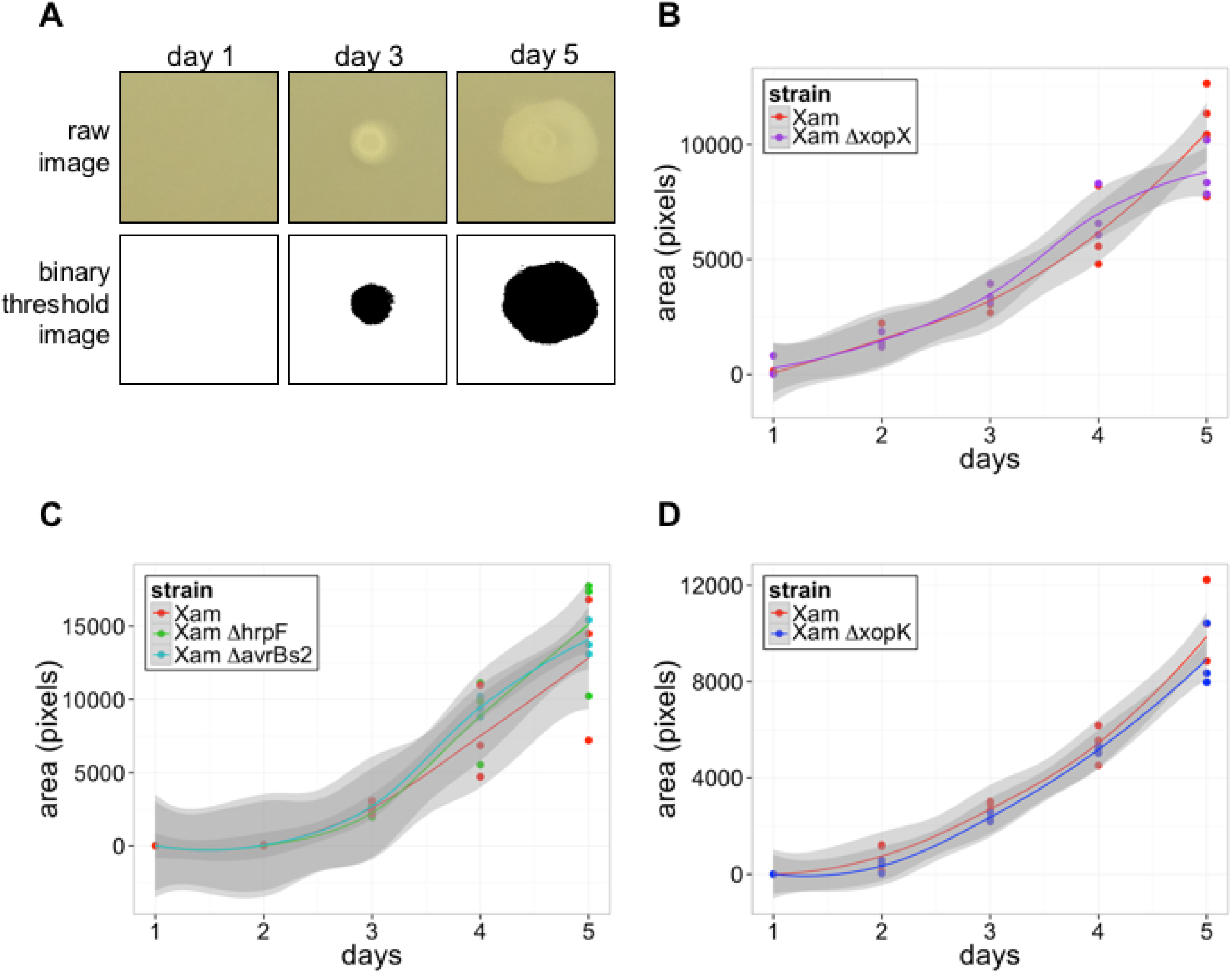
*In vitro* motility of *Xam* T3E mutants. Bacteria were spotted in the center of plates containing NYG media with 0.25% agar. Images were taken of the plates over several days, and the area of bacterial spread was quantified using ImageJ. A, Example of binary thresholding used to select the area of bacterial spread from the images. B-D, Quantification of bacterial spread for the *Xam* T3E mutants, relative to the wild-type strain. Dots represent individual data points, and local regression fitted curves are plotted for each bacterial strain. Shaded areas represent the 95% confidence interval for each strain.

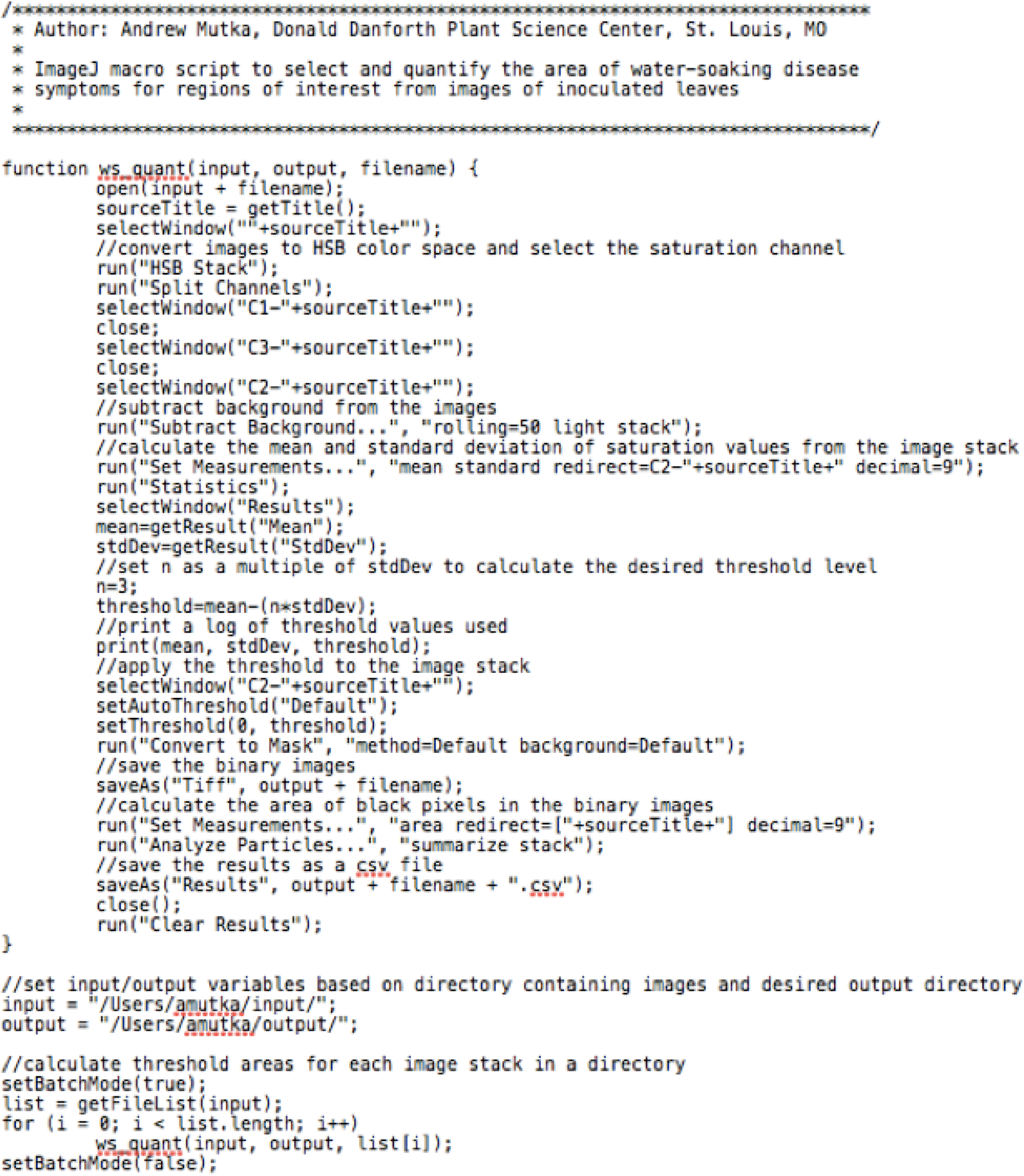

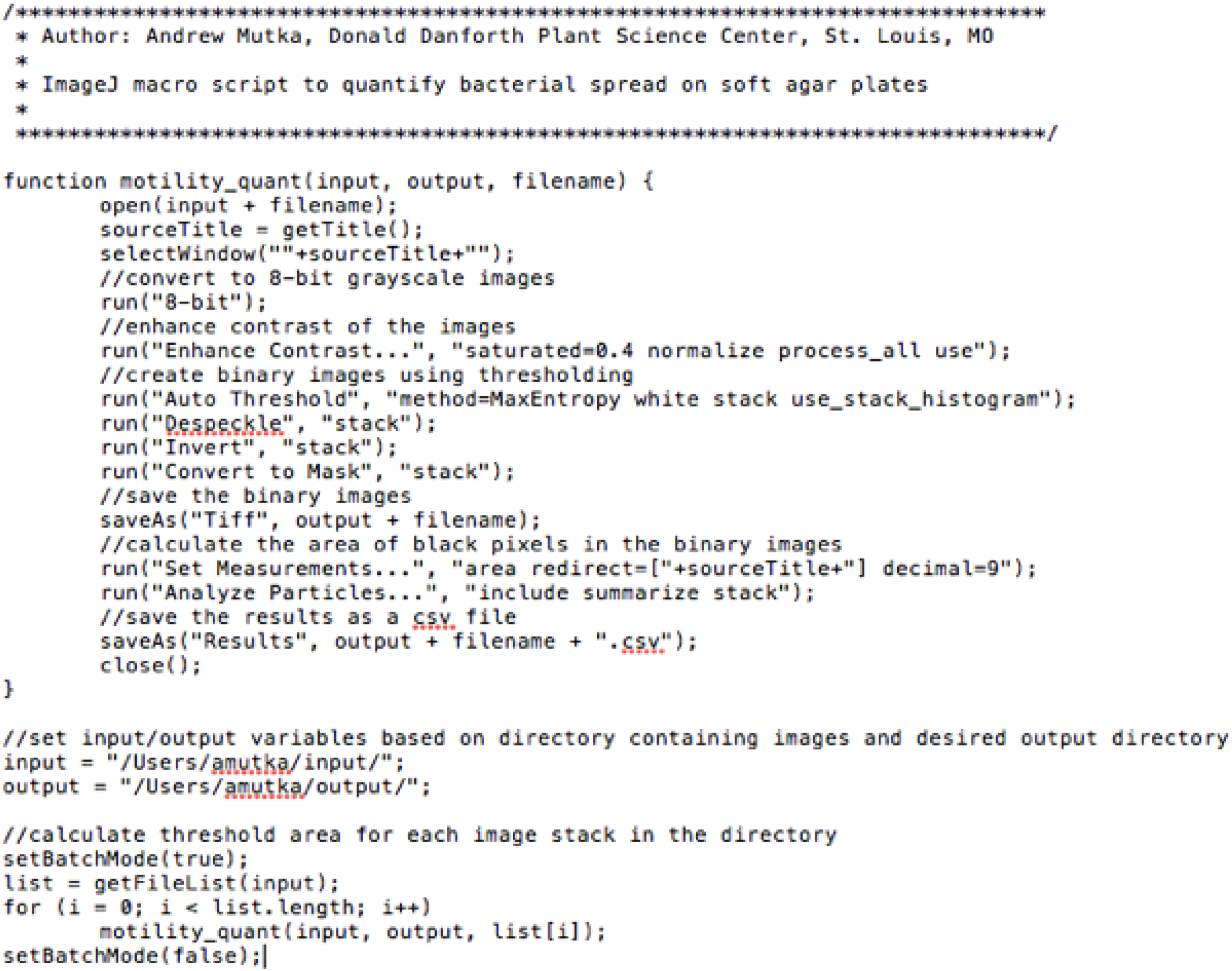

